# Application of the Mesolens for sub-cellular resolution imaging of intact larval and whole adult *Drosophila*

**DOI:** 10.1101/267823

**Authors:** Gail McConnell, William B. Amos

**Author notes:** Corresponding author’s: email address. Corresponding author’s.

## Abstract

In a previous paper (McConnell et al., 2016) we showed a new giant lens called the Mesolens and presented performance data and images from whole fixed and intact fluorescently-stained 12.5-day old mouse embryos. Here we show that using the Mesolens we can image an entire *Drosophila* larva or adult fly in confocal epifluorescence and show sub-cellular detail in all tissues. By taking several hundreds of optical sections through the entire volume of the specimen, we show cells and nuclear details within the gut, brain, salivary glands and reproductive system that normally require dissection for study. Organs are imaged *in situ* in correct 3D arrangement. Imaginal disks are imaged in mature larvae and it proved possible to image pachytene chromosomes in cells within ovarian follicles in intact female flies. Methods for fixing, staining and clearing are given.

## 1. INTRODUCTION

*Drosophila* has been described as ‘too small for easy handling but too large for microscopy’ (Chyb and Gompel, 2013). The Mesolens, with its unusual combination of low magnification and high numerical aperture, solves the size problem without compromising image resolution (McConnell *et al.*, 2016), but the impermeability of the cuticle prevents the use of many preparative methods. Dissection overcomes the second problem and has allowed the observations, including those with photoproteins or hybridization probes at the highest resolution of a conventional microscope (Singh *et al.*, 2011; Long *et al.*, 2017; Nern *et al.*, 2015), but provides no spatial information about the original relationship of the dissected structures. Paraffin sectioning is slow and often fails to preserve antigens and fine structure (Demerec, 2008). Light-sheet illumination has been successful in imaging specimens up to 400 μm long (Tomer *et al.*, 2012). Other methods such as micro-CT (Matsuyama *et al.*, 2015) and optical coherence tomography (McGurk *et al.*, 2007) provide only low-resolution images.

We have here taken a different approach, using the power of our novel lens system (McConnell *et al.*, 2016) to capture detail at high resolution throughout a volume of over one hundred cubic millimetres: large enough to image three or more mature *Drosophila* larvae or adults at once.

## 2. MATERIALS AND METHODS

### 2.1 Fly husbandry and stocks

Flies were reared on ‘normal’ laboratory food (1 L recipe: 80 g corn flour, 20 g glucose, 40 g sugar, 15 g yeast extract, 4 ml propionic acid, 5 ml p-hydroxybenzoic acid methyl ester in ethanol, 5 ml ortho butyric acid) at room temperature under 12 h/12 h light/dark conditions.

### 2.2 Specimen preparation

A classic non-aldehyde fixative (ethanol:acetic acid, 3:1 by volume) was employed because of its known rapid penetration and good preservation of chromatin (Baker, 1958). Fixation in 4% paraformaldehyde gave similar results to the use of ethanol:acetic, albeit with a slight reduction in fluorescence signal. The stains chosen, propidium iodide (PI) and HCS CellMask Green, were found to give a result similar to the haematoxylin and eosin of conventional histology but were more suited to confocal microscopy because of their fluorescence. By piercing the cuticle of chilled flies, as recommended by Bodenstein (Bodenstein, 1950), the ingress of fixative and subsequent processing fluids was facilitated.

Wild type *Drosophila melanogaster* (3^rd^ instar larvae and imago) were anaesthetised by placing in a plastic tube surrounded by dry ice for 2 minutes. When chilled to near-immobility, each specimen was lifted using fine tip forceps and placed into a 66 mm diameter glass Petri dish under a stereo microscope. The fly or larva was held in place with the same forceps and a needle made by electrolytic sharpening (Brady, 1965) of tungsten wire 0.5 mm in diameter was used to make two punctures in the abdomen and thorax of the adult, and the body of the larva, allowing the ingress of fixative, bleach, stains and clearing solutions.

Once punctured, the specimens were fixed in 3:1 by volume ethanol/acetic acid (E/0650DF/17, Fisher Scientific & 695092-2.5L, Sigma-Aldrich) by placing the specimens in a small Petri dish of fixative mixture on a gentle rocker for 3 hours at room temperature. Following fixation the specimens were washed three times in PBS (10010023, ThermoFisher Scientific), 5 minutes each. To bleach and to aid subsequent tissue clearing the specimens were placed in 35% H_2_O_2_ (349887-500ml, Sigma-Aldrich) for 18 hours, again with gentle rocking. Specimens were removed from 35% H_2_O_2_ and washed in PBS (3x, 5 minutes). Next, the bleached specimens were treated with 100 μg/ml RNAse (EN0531, ThermoFisher Scientific) in PBS for 1 hour at room temperature with gentle agitation. Without washing, the specimens were then added to 10 μM propidium iodide (P4864-10ml, ThermoFisher Scientific) and gently rocked for 4 hours. For two-colour staining HCS Cellmask Green, 20 μL of a 10 mg/ml stock solution of HCS Cellmask Green (H32714, ThermoFisher Scientific) was added to 10 ml PBS, and this was applied at the same time and for the same duration as the PI stain. After the dye loading and for subsequent steps, the specimens were covered with aluminium foil to reduce bleaching by ambient light.

The fluorescently-stained specimens were washed in PBS (3x, 5 minutes) and for ease of handling the specimens were mounted in small blocks (8 mm diameter) of 0.8% agarose (05066-50G, Sigma-Aldrich). The agarose-mounted fluorescent specimens were then dehydrated through a methanol series (50% MeOH, 75% MeOH, 100% anhydrous MeOH, 100% anhydrous MeOH, each step 1 hour) with gentle agitation. BABB was introduced by placing the specimens in a 1:1 by volume mixture of anhydrous MeOH (322415-1L, Sigma-Aldrich) and BABB, the latter being a 1:2 mixture of benzyl alcohol (402834-500ml, Sigma-Aldrich) to benzyl benzoate (B6630-1L, Sigma-Aldrich) and rocked for one hour. It is noted that glass dishes are essential at this stage because BABB dissolves plastic. The specimens were removed from the MeOH/BABB mix and placed in 100% BABB and gently rocked for at least 24 hours before imaging.

### 2.3 Specimen mounting procedure

Each specimen was placed in a holder constructed by cementing an aluminium spacer plate to a larger-than-standard microscope slide using a proprietary adhesive (UHU MAX) to create a leak-proof seal resistant to immersion oil. The slide was type1529100092 (Marienfeld), measuring 100 × 76 × 1 mm and the aluminium plate measured 80 × 70 × 3 mm with a central hole 10mm in diameter. BABB was added to cover the specimen and a large coverslip placed on top (70 mm × 70 mm) type 1.5 0107999098 (Marienfeld) avoiding bubbles. The Mesolens was used with oil immersion and capillarity proved insufficient to preserve the oil column (up to 3 mm high) between lens and coverslip for long periods. A special chamber was therefore constructed to allow a metal ring with a nitrile ‘O’ ring 30mm in diameter on its underside to be held against the coverslip. A computer-aided design drawing of the specimen holder and chamber to support long-term immersion is shown in Figure 1. After assembly, immersion oil was added, creating a stable oil bath within the ring with the coverslip as its base and the front of the Mesolens dipping into it.

**Fig 1.**
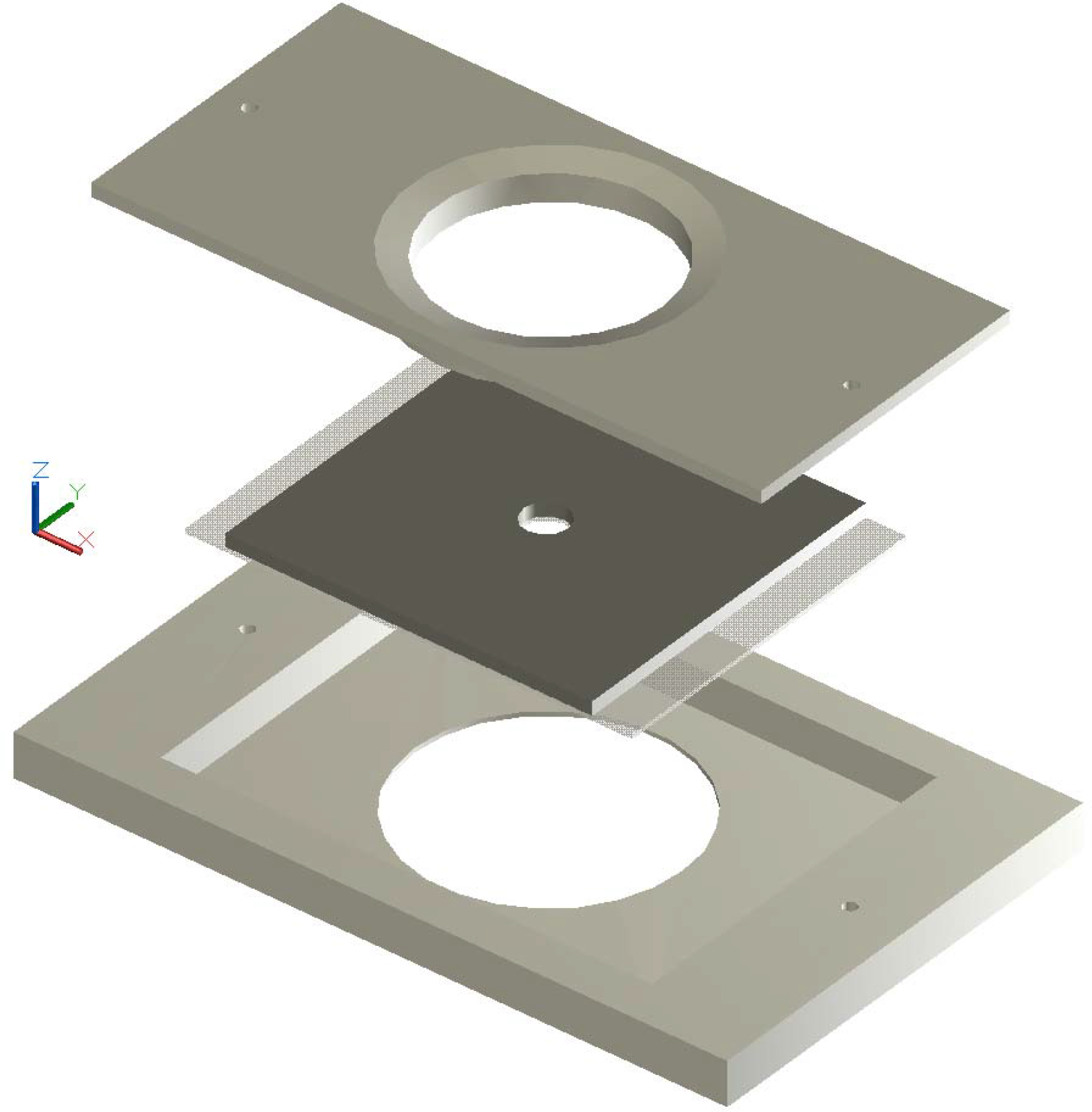
A computer-aided design drawing of the disassembled specimen holder and multi-part chamber to support long-term use of immersion fluid for imaging *Drosophiila* specimens with the Mesolens. The top and bottom sections correspond to the immersion chamber, while the specimen is mounted on a specially-large slide under a coverslip with an aluminium spacer between slide and coverslip in the mid-section. The top plate includes a 30 mm diameter ‘O’ ring (not shown) that is brought into contact with the coverslip on top of the 100 mm long specimen slide, and the bottom plate creates a stable base. Two screws (not shown, though through holes are presented) bring the three sections together. Immersion fluid is added to the bath created by the contact of the top plate and coverslip of the specimen slide for long-term imaging.

### 2.4 Imaging conditions

Details of the Mesolens are already reported (McConnell *et al.*, 2016), so only the imaging parameters used in these experiments are described here. For fluorescence excitation of the HCS Cellmask Green and PI stains, laser powers of no more than 3 mW and 5 mW (Laserbank, Cairn Research) at wavelengths of 488 nm and 561 nm were used for simultaneous dual-wavelength excitation and detection, with less than 200 μW of total laser power incident on the specimen during scanned imaging. Fluorescence from the HCS Cellmask Green stain was spectrally separated from the 488nm excitation using a 550 nm dichroic filter (DMLP550R, Thorlabs) and a 525/39 nm bandpass filter (MF525-39, Thorlabs) before detection with a photomultiplier tube (P30-01, Senstech) Similarly, the red fluorescence from the PI stain was separated from the yellow excitation using the same dichroic filter with a 600 nm long-wave pass filter (FEL0600, Thorlabs), and was detected using a second photomultiplier tube (P30-09, Senstech). A galvo mirror scan speed of 40 Hz was used to image all specimens. Though the imaging speed is slow (approximately 50 seconds per image), the point-scanning confocal method supports optical sectioning to minimise out-of-focus fluorescence. We also note that even if the out-of-focus fluorescence could be tolerated for very thin or sparsely labelled specimens, for full Nyquist sampling a camera chip with over 200 Megapixels would be required. At present, to the best of our knowledge, this chip technology is not available in a commercial scientific camera product.

For all experiments, we chose the frame size based on the size of the specimen and, because the Mesolens is not a zoom lens, the numerical aperture of the lens does not change with frame size. As such, there is no resolution improvement to be gained by increasing the frame size. Instead the number of pixels in each frame was set to exceed the Nyquist sampling limit.

## 3. RESULTS

Anatomical identification was by reference to Demerec (Demerec, 2008).

### 3.1 Internal structure of the larva

In the 3^rd^ instar larva (Fig 2A) there is almost no free space: cellular structure, often of the net-like cytoplasm of fat-body cells, filled the entire internal volume. As expected, green fluorescence of cytoplasm produced by HCS CellMask Green and red fluorescence of nuclei and chromosomes due to propidium iodide (PI) were observed. Unstained specimens showed only a weak blue/green autofluorescence. PI also seemed to stain cuticular spicules on the exterior. There was little sign of damage due to the piercing of the exoskeleton. In a typical imaging session, up to 220 confocal optical sections were obtained in the 850 μm thickness of the larva and adjacent sections showed quite different patterns of nuclei and even of cells, since the optical section thickness was less than 4 μm. The time taken to image each plane with two-channel detection was 50 seconds. It proved unnecessary to increase laser power to obtain sufficient fluorescence from deeper sections.

**Fig 2.**
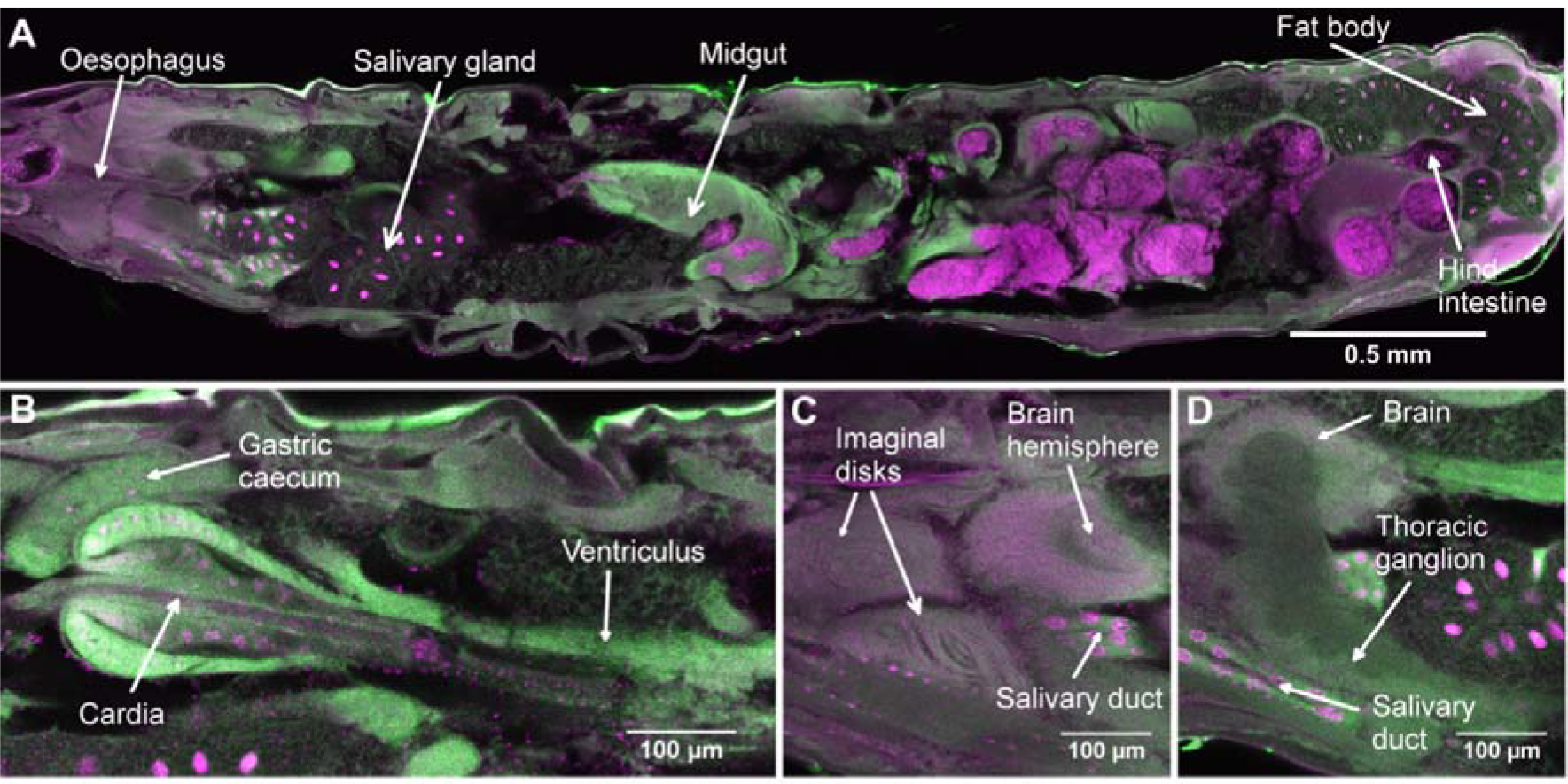
Confocal optical sections of an intact 3^rd^ instar larva of *Drosophila*. 2A is a median sagittal section passing through the oesophagus. The field size of 2A is 4 mm × 0.92 mm (8000 pixels by 1904 pixels), with a pixel size of 0.5 μm in both the x and y dimensions and 3.99 μm in the z dimension. Fluorescence of propidium iodide is shown in magenta and is localized in nuclei, gut contents and some cuticular spines. HCS CellMask Green, intended as a cytoplasmic label, is shown in green, and highlights tissues with dense cytoplasm and stains the exoskeleton also. 2B is another section at a different level (83.79 μm deeper into the specimen), passing through a gastric caecum and the cardia. 2C, which is 95.76 μm closer to the near surface of the specimen than 2A, shows a dorsal brain hemisphere and the imaginal disks reveal a concentric multi-layered structure and primordial optic ganglia of the brain, with the pinkish hue of the cortex due to the massed nuclei and the darker fibrous core. 2D, which is between 2A and 2C (31.92 μm closer to the near surface than 2A), shows the thoracic ganglion and salivary gland relative to the brain hemisphere. This dataset is representative of n=8 larvae imaged using the same method.

The cuticle of the larva was predominantly stained with HCS CellMask Green, as were the peripheral muscles, though the striations were not made visible by this stain. Some expected PI staining was absent e.g. the large (10-15 μm diameter) epidermal nuclei were seen only in restricted regions of the larva. It was possible to trace the alimentary canal completely from mouth to anus. The oesophagus (Fig 2A) showed as a green-fluorescing tube passing through the brain and continuous with the central canal of the cardia (Fig 2B, 2C). Nuclei in the gastric caeca were clearly visible. The massive and convoluted midgut also took up the green basophilic stain intensely, and the gut contents were red-fluorescent, probably because of the DNA content of yeast in the food. The hind-intestine (Fig 2A) showed a bright green fluorescence in its thick wall, which had the appearance of an internally-toothed ring in cross-section.

The brain hemispheres, with their fibrous centres and outer cellular layers, were clear (Fig 2C), and the segmentation of the ventral ganglia was visible by virtue of the scalloped boundary between cortex and inner core (Demerec, 2008). Several imaginal disks were revealed as globular bodies, with characteristic folded concentric layers (Fig 2C). The nuclei were barely visible.

The large polytene nuclei of the salivary glands were clear (Fig 2A). Individual polytene chromosomes could be seen, particularly in the nuclei of the salivary duct. The distal portion of each salivary gland, was strongly stained with HCS CellMask Green (Fig 2D). The fat cells also showed prominent polytene nuclei suspended at the centre of a net-like remnant of the cytoplasm, from which the fat globules had been extracted during fixation.

### 3.2 Internal structure of the adult fly

A quick overview of the imaging potential of the Mesolens can be gained by viewing Movie 1, discussed below.

Fig 3 shows a projected image of the entire volume of a female fly obtained from a confocal image stack, and shows how fine detail is visible at all depths. There was little background fluorescence because the cytoplasmic stain was not used in this case: only PI was used.

**Fig 3.**
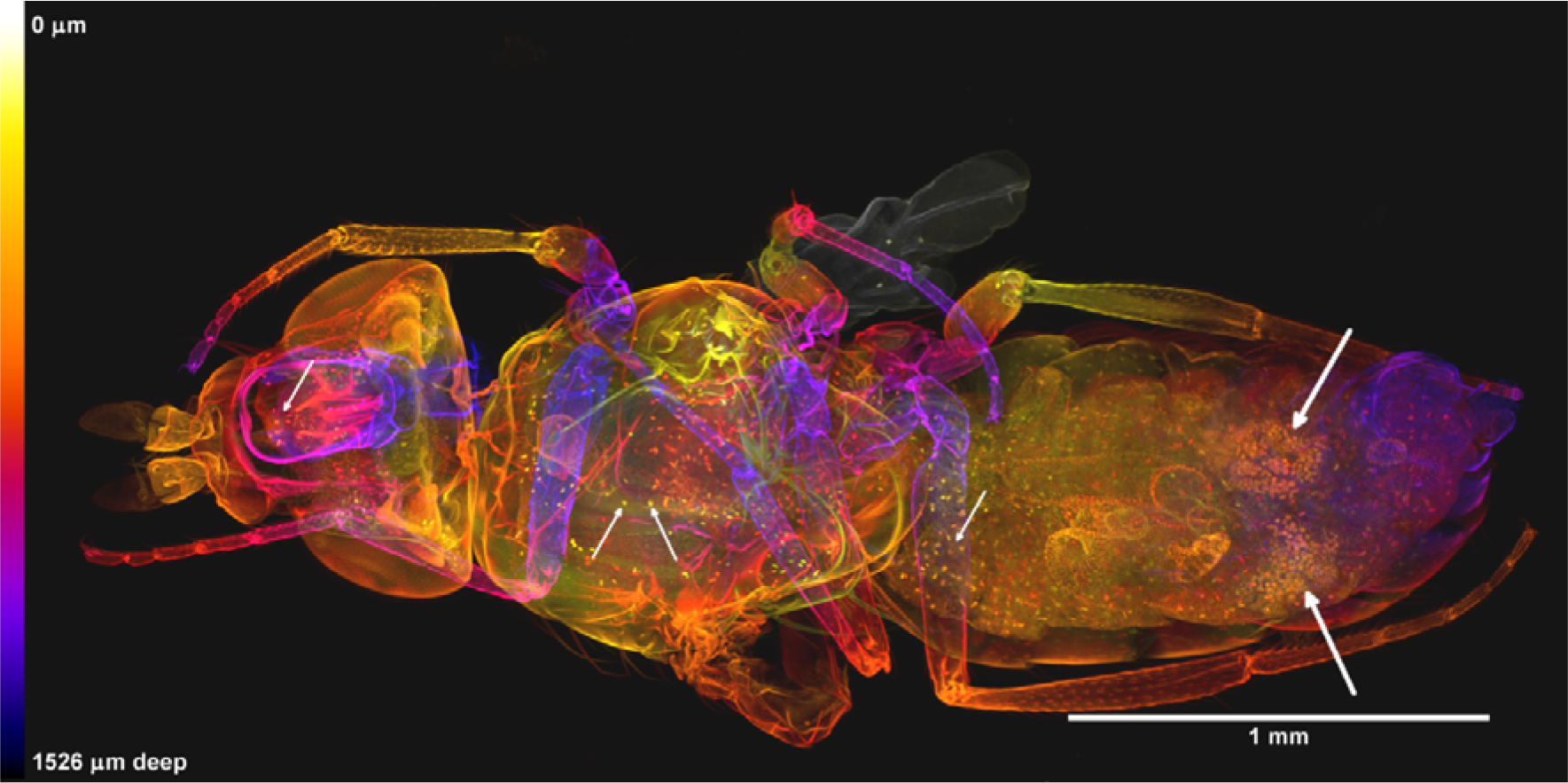
Whole female *Drosophila* newly emerged imago in dorsal view. This image is composed by projection of 242 optical sections taken with an axial separation of 6.3 μm, forming a z-stack 1.53 mm deep. The sections are colour-coded for depth according to the scale shown, in which near sections are yellow and the far ones purple or dark grey. Only propidium iodide was used as a fluorochrome, revealing both cuticle and nuclei of individual cells in the interior. The orange clusters of nuclei in the abdomen are those of the ovaries (indicated with large white arrows at the right hand side of the image). The smaller white arrows on the left of the image indicate PI-positive bodies that may be parasites. This dataset is representative of n=7 adult flies imaged using the same method.

Fig 4A shows a different female fly, stained with both HCS CellMask Green and PI. In the head, the three ocelli (not all shown in the figure), revealed the form of their lenses and the cellular structure of the internal strand (peduncle) linking them to the brain, as shown in Fig 4D. The head capsule was stained strongly with PI. The brain was visible as a HCS CellMask Green positive body. Within each ommatidium of the compound eye the discrete group of retinula cell nuclei was clear (Fig 4B, with a zoomed region of 3B shown in 3C, which reveals the fine detail of the cell nuclei), as was a deeper layer of ganglion cell nuclei lying within each optic lobe. The antennal muscles and nerves stained strongly with HCS Cellmask Green (Fig 4A). Detail of the intricate mouthparts included chitinous structures such as the rostrum taking up PI and the extensors of the labellum and the pharynx, the green-fluorescing stain.

**Fig 4.**
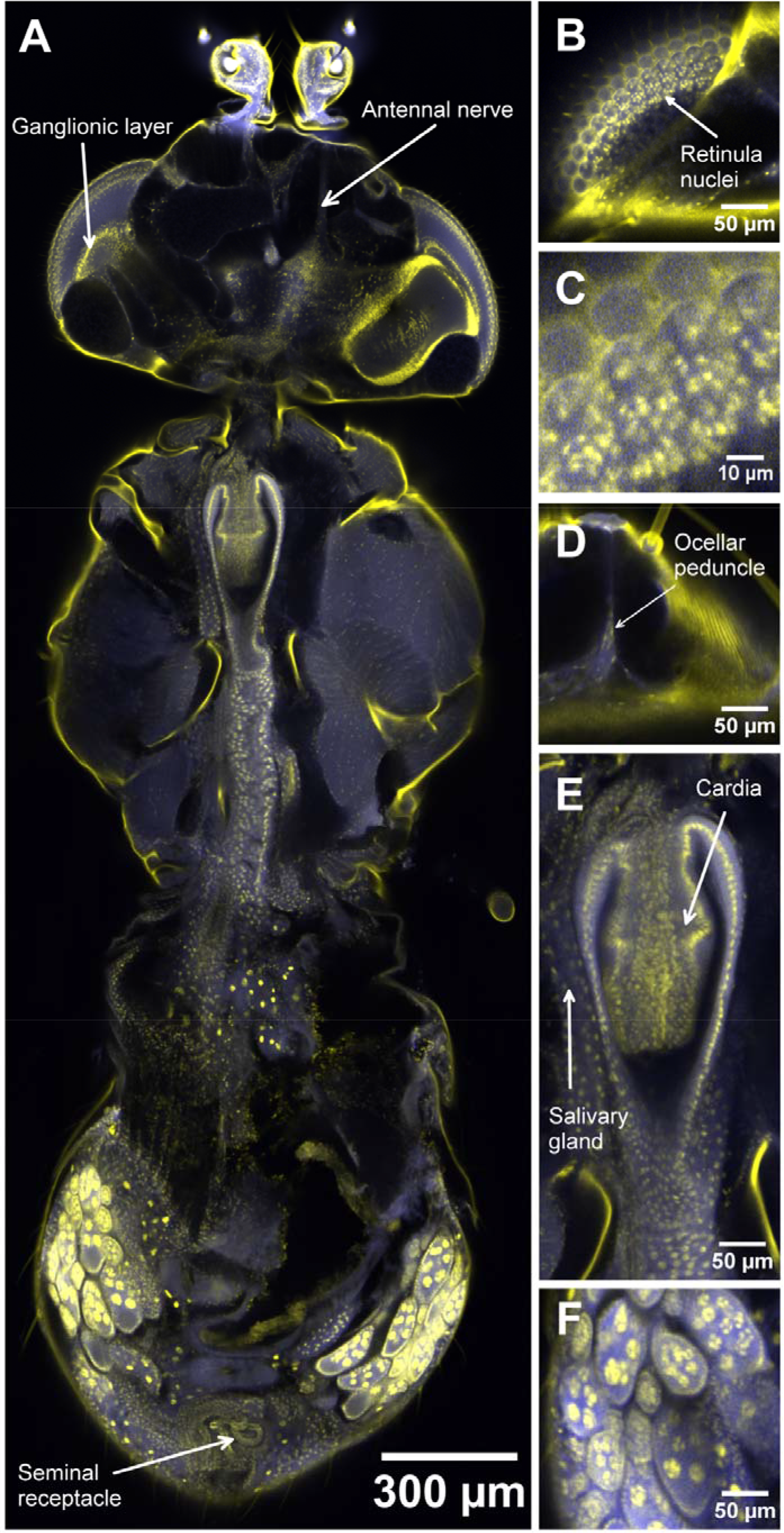
Confocal optical section and digitally zoomed regions of a whole adult female *Drosophila.* 4A is an optical section of an intact female *Drosophila* imago, mounted to present a dorsal aspect with the focal plane approximately midway between dorsal and ventral surfaces, at a depth of 892.11 μm into the specimen. The field size of 4A is 2.934 mm × 1.349 mm (11736 pixels by 5396 pixels), with a pixel size of 250 nm in both the x and y dimensions and 3.67 μm in the z dimension. The fluorescence of propidium iodide is here shown yellow and is localized in the exoskeleton, particularly in the thorax, and in nuclei, which results in epithelia appearing as lines of yellow dots in transverse section. The strongest propidium staining is in the left and right ovaries, of which the epithelia are distinct and the pachytene chromosomes are stained very strongly in the mitotic and meiotic divisions of oogenesis. Yellow nuclear zones are also visible in the ganglionic layer of the left compound eye and around the surface of the optic region of the brain sectioned on the right, and in the wall of the tubular seminal receptacle. Lines of nuclei are also visible in the thoracic flight muscles, in which the cytoplasmic stain HCS Cellmask Green provides a blue background. 4B, obtained at a depth of 613.08 μm into the specimen, is an enlarged detail of a glancing section passing through the compound eye, in which the groups of retinula cell nuclei are visible. Fig 4C shows a software zoomed version of Fig 4B, revealing the retinula nuclei fine detail (we note that Fig 4C is presented with a 10 μm scale bar, while Figs 4B, and 4D-F are presented with a 50 μm scale bar). Fig 4D, obtained at the same focal plane as 4B, shows the ocellar peduncle. Fig 4E shows detail of the infolded epithelia of the cardia and the posterior extension of the gut from it, imaged at a depth of 903.9 μm. Several ovarian follicles are visible in 4F, obtained at an imaging depth of 994.29 μm, with chromosomes in nurse cells, each follicle surrounded by epithelium. This dataset is representative of n=7 adult flies imaged using the same method.

In the thorax the flight muscle fibres were strongly stained with HCS CellMask Green, and their lines of nuclei with PI. As in the larva, striations in the muscle fibre were not visible. We believe this to be a consequence of the staining method, rather than the resolution of the image. In the ventral part the connection of the oesophagus to the cardia and the cell layers of the cardia and ventriculus (Figs 4A and 4E) and the tubular salivary glands running alongside were shown in great detail. Ventral to this the thoracic ganglionic mass, constricted into three segments by the exoskeleton, lay above the ventral muscles.

In the abdomen (Fig 4A), the gut, Malpighian tubules and reproductive system, were all shown in sub-cellular detail. The crop and ventriculus could be seen and their lumina could be traced through to the rectal sac with its papillae. The Malpighian tubules were distinguished clearly by their large and intensely-stained nuclei.

In the male, stages of nuclear transformation during sperm formation could be seen (data not shown here) and in females the ovaries were obvious, with metaphases of meiotic and mitotic divisions during oogenesis being clearly visible by the dense staining of chromosomes or bivalents with PI in the mature follicles Figs 4A and 4F. The distal regions of the female system were well shown, with the coiled seminal receptacle and the paired spermathecae (Fig 4A) and their ducts visible. Some confocal optical sections (not shown) showed the uterus greatly distended by a single egg, with uterine cells and nuclei visible, and the yolk of the egg shrunken and irregularly-shaped in fixation.

With PI alone the exoskeleton was the brightest structure. The rotating display, shown in Movie 1 was made by constructing a series of projections at different angles (2 degrees, through 360 degrees of rotation) using a maximum brightness projection algorithm in Icy (de Chaumont et al., 2012), which effectively eliminated all but the exoskeletal signal. However, colour depth-coding of the same data (Fig 3) did not eliminate the weaker staining and showed interior structures such as nuclei. Fig 3 also shows numerous small elongated or fusiform PI-positive bodies distributed through many tissues which may perhaps be spores of a microsporidian parasite (Franzen *et al.*, 2005).

## 4. DISCUSSION

These results show that our approach may be a useful additional tool for *Drosophila* research, not replacing dissection and high-resolution observation of individual organs but allowing a whole-body examination without danger of loss of cells during dissection and to see the intricate spatial relationship between parts, including clones of cells, during development. It is also far preferable to dissection for finding small objects within the large volume of the insect body.

Unfortunately, methanol dehydration quenches the fluorescence of photoproteins, but recent work suggests that this problem may be overcome by the use of strongly alkaline buffers during dehydration in ethanol before immersion in BABB (Schwarz *et al.*, 2015). It is also likely that other clearing solutions could be substituted for BABB (Richardson and Lichtmann, 2015), and that clearing of live larvae may be possible using the refractive-index tunable and non-toxic clearing method recently by Boothe et al (Boothe *et al.*, 2017). Unlike mammalian tissue (Hama et al., 2011; Ke, Fujimoto and Imai, 2013) we did not observe significant tissue shrinkage using the BABB clearing method.

Our finding that small PI-positive bodies, probably pathogenic organisms, could be imaged in all tissues of certain adult flies suggests that our methods could be applied in the study of infection, immunity and parasitism in *Drosophila*. The alimentary canal of *Drosophila* is of interest as a model of bacterial infection, e.g. (Ekström and Hultmark, 2016), and the ability shown here of imaging the entire gut contents may prove useful.

## CONCLUSION

The chief conclusion of this work is that structures within the body of adult and larval forms of an insect are amenable to high-resolution optical microscopy and that the entire body may be viewed in single confocal images using a Mesolens. We offer this demonstration in the hope that this approach may facilitate studies of the distribution of clones of cells and of other phenomena that involve the entire organism. We hope that the present results will prove of interest to researchers using *Drosophila* as a model of human disease, or wherever a global view of the entire organism is needed, with sub-cellular resolution. We are currently extending this work to *Culex*, *Tribolium* and other intensively-studied insects.

## Acknowledgements

The authors would like to thank Lince José Casal and Peter Lawrence (Department of Zoology, University of Cambridge) for helpful discussions and advice on the manuscript. We also thank Lee McCann (Department of Physics, University of Strathclyde) for his help in the preparation of Fig. 1.

## Data availability

All datasets supporting this work are available at https://strathcloud.sharefile.eu/d-s8c09b9e95b143679.

## Movie 1

Z-series and rotating 3D view of whole female *Drosophila* imago. The first part of the movie shows a z-series of 242 images through a whole female Drosophila imago, with all external and internal structure visible. The second part of the movie shows mainly the exterior of the fly in a rotating 3D reconstruction of the same specimen, created using the ‘3D Rotation’ plugin in the image processing software Icy. A series of projections at different angles (2 degrees, through 360 degrees) was made using a maximum brightness projection algorithm, which effectively eliminated all but the bright exoskeletal signal. We note that unlike projections of z-series made with conventional lenses, these projections do not show blurring in the optically axial direction (here used as the rotation axis). This is because the axial resolution length is small compared with the height of the specimen.

